# Integrating telemetry data into spatial capture-recapture modifies inferences on multi-scale resource selection

**DOI:** 10.1101/131144

**Authors:** Daniel W. Linden, Alexej P. K. Sirén, Peter J. Pekins

## Abstract

Estimating population size and resource selection functions (RSFs) are common approaches in applied ecology for addressing wildlife conservation and management objectives. Traditionally such approaches have been undertaken separately with different sources of data. Spatial capture-recapture (SCR) provides a framework for jointly estimating density and multi-scale resource selection, and data integration techniques provide opportunities for improving inferences from SCR models. Here we illustrate an application of integrated SCR-RSF modeling to a population of American marten (*Martes americana*) in alpine forests of northern New England. Spatial encounter data from camera traps were combined with telemetry locations from radio-collared individuals to examine how density and space use varied with spatial environmental features. We compared multi-model inferences between the integrated SCR-RSF model with telemetry and a standard SCR model with no telemetry. The integrated SCR-RSF model supported more complex relationships with spatial variation in third-order resource selection (i.e., individual space use), including selection for areas with shorter distances to mixed coniferous forest and rugged terrain. Both models indicated increased second-order selection (i.e., density) for areas close to mixed coniferous forest, while the integrated SCR-RSF model had a lower effect size due to modulation from spatial variability in space use. Our application of the integrated SCR-RSF model illustrates the improved inferences from spatial encounter data that can be achieved from integrating auxiliary telemetry data. Integrated modeling allows ecologists to join empirical data to ecological theory using a robust quantitative framework to better address conservation and management objectives.

## Introduction

Understanding the environmental features that influence variation in species abundance or density is a common objective in wildlife conservation and management. Approaches to estimating population size and habitat selection have traditionally required distinct forms of data collection and separate modeling methods; spatial capture-recapture (SCR) allows for joint estimation of both ecological processes using a single data source or through integration of multiple data sources in a single analytical framework (Royle, Fuller & Sutherland 2017). The development of SCR was initially motivated by the need to address the spatial dynamics of field sampling and animal movement to improve density estimation from capture-recapture methods (Efford 2004; Royle & Young 2008). By using spatial information on the location of observations, SCR combines a point process model for the distribution of individuals in a population with a probability model for the encounters or captures of those individuals. As with other recent statistical advances in ecology (Gimenez *et al.* 2014), the hierarchical model formulation of SCR has enabled custom data integration techniques and, therefore, expanded the scope of inferences possible from trapping and other spatial encounter data (Sollmann *et al.* 2013a; Chandler & Clark 2014). While these developments are promising, as the complexity of modeling methods increases so do the data demands for parameter estimation (Auger-Methe *et* al. 2016) and the need for practical implementation options (Bolker *et al.* 2013). Producing valid and useful inferences for helping achieve objectives in applied ecology requires finding a balance between comprehensive models and logistically feasible data.

Royle *et al.* (2013) developed a spatial capture-recapture model that integrates telemetry information and resource selection functions (RSFs) to provide improved accuracy and precision for density estimation. The improved accuracy is particularly relevant when resource selection at one or more spatial scales affects the distribution of individuals (i.e., second-order selection; Johnson 1980) and/or individual space use (i.e., third-order selection). While the approach seems promising and uses commonly collected wildlife data, it has been rarely applied in the literature. Proffitt *et al.* (2015) included a resource selection function in an SCR model of a large carnivore but estimated the function with telemetry data first before using the predictions as a covariate in their SCR model fitting. They implied that the Royle *et al.* (2013) model represented a “methodologically intensive joint estimation framework” which might preclude interested users from easily applying it. The multi-step approach to RSF integration in SCR is a potentially interesting and practical compromise, whereby a complex function is reduced to a univariate prediction to serve as a single spatial covariate in the model (Efford 2015). Proffitt *et al*. (2015) used this method to address spatial variation in density – they did not incorporate it in their encounter probability model, despite having data that may have allowed for modeling variation in individual space use. In general, SCR models are more sensitive to the structure of the encounter model than the density model, as the latter has been found to be highly flexible to misspecification (Efford & Fewster 2013; Royle *et al.* 2014) while the former has long been a focus of refinements to capture-recapture methods (Dorazio & Royle 2003). As referenced earlier, individual heterogeneity to capture was a primary motivation for SCR (Efford 2004). Thus, in SCR applications where individual space use is hypothesized to be highly variable, accounting for third-order selection in the encounter model may be important regardless of whether ancillary telemetry data are available.

The task of estimating variation in space use with a spatial capture-recapture design alone is made difficult by the required sample sizes in the various data dimensions. Importantly, the number and configuration of trapping devices needs to be such that adequate coverage across some gradient of habitat resources is achieved. This requires some traps to be placed within or adjacent to relatively poor habitat, even though doing so may result in little information (i.e., zero encounters) being collected for the logistical effort expended. Since the placement of traps will interact with both the distribution of individuals on the landscape and individual space use, the number of encounters at a trap will be a function of the realized dynamics of both processes (Royle *et al.* 2014). Thus, an optimal trap layout for estimating multi-scale resource selection will purposefully expend trapping effort in locations where few individuals exist and where individuals, even when present nearby, may be unlikely to visit. Such a design creates tension with the general goal of obtaining as many encounters of as many individuals as possible to enable model fitting and accurate parameter estimation (Royle *et al.* 2014).

Here, we illustrate an application of the integrated SCR-RSF model developed by Royle *et al.* (2013) using a multi-year study on American marten (*Martes americana*) in New Hampshire, USA (Sirén *et al.* 2016a; Sirén *et al.* 2016b). Marten are a forest-sensitive meso-carnivore often used as an indicator species for forest conservation and management given their vulnerability to anthropogenic disturbance and climate change (Carroll 2007). We used remote camera stations to photograph and identify individual marten according to their unique pelage patterns (Sirén *et al.* 2016a) and combined these data with telemetry locations collected on a subset of radio-collared individuals (Sirén *et al.* 2016b). The sampling occurred across a heterogeneous alpine forest landscape that was recently modified by a wind farm installment and within which marten were shown to be differentially selecting resources at multiple scales. Using the integrated SCR-RSF likelihood (Royle *et al.* 2013), we estimated marten density and parameters associated with multi-scale resource selection and compared the resulting multi-model inferences both with and without the telemetry data integration. Notably, we modified the likelihood to accommodate a lack of independence between the data sources, given that all radio-collared individuals were also photo-captured by camera traps. This integrated SCR-RSF model was added to the R package oSCR (Sutherland, Royle & Linden 2016) to facilitate use by interested researchers. Our example here provides further evidence that improved inferences on variation in population density are possible when additional information on movement and resource selection from telemetry data are integrated with spatial capture-recapture models.

## Materials and methods

### STUDY AREA AND SAMPLING

We sampled marten in a ∼62 km^2^ area of New England-Acadian forest in northern New Hampshire, USA during 2010–2012 (Figure S1). Extensive descriptions of the study area and sampling details have been previously outlined (Sirén *et al.* 2016a; Sirén *et al.* 2016b). The area was centered on some mountains containing a new 33-turbine wind farm with elevation ranging 624–1045 m. Forests were primarily mature conifers at high elevations (>823 m) and a mix of mature and commercially harvested hardwoods at lower elevations. Topography was variable with rugged terrain dispersed throughout and winters in the region (1948–2013) were cold with high annual snowfall (average = 288 cm; range = 79–881 cm; National Climate Data Center: http://www.ncdc.noaa.gov).

Marten were photo-captured by remote cameras during 2 winters (Sirén *et al.* 2016a) and live-captured year-round to maintain 6–10 individuals with active VHF radio collars during the study (Sirén *et al.* 2016b). Telemetry fixes from collared individuals were obtained weekly using standard methods of triangulation; here, we restricted the data to fixes collected during the 2 leaf-off (i.e., winter) time periods (14 Nov 2010 to 15 May 2011 and 16 Oct 2011 to 15 May 2012) which overlapped with the winter sampling by remote cameras (14 Feb 2011 to 2 Apr 2011 and 3 Jan 2012 to 19 Jan 2012). Location error was estimated to be 2 ha (Sirén *et al.* 2016b). We used 30 total camera trapping stations during each sampling period that were baited with sardines and a commercial skunk (*Mephitis mephitis*) lure. Stations were operational for 12 d in 2011 and 8 d in 2012, with bait replaced halfway through, and were located in a nonrandom pattern to achieve a spacing of 500–950 m (Figure 1). Marten photo-captures were identifiable to individual when ventral patches were clearly visible and multiple observers could agree on assignment; sex was assigned to un-collared individuals based on morphological comparisons with collared individuals (Sirén *et al.* 2016a).

**Figure 1.**
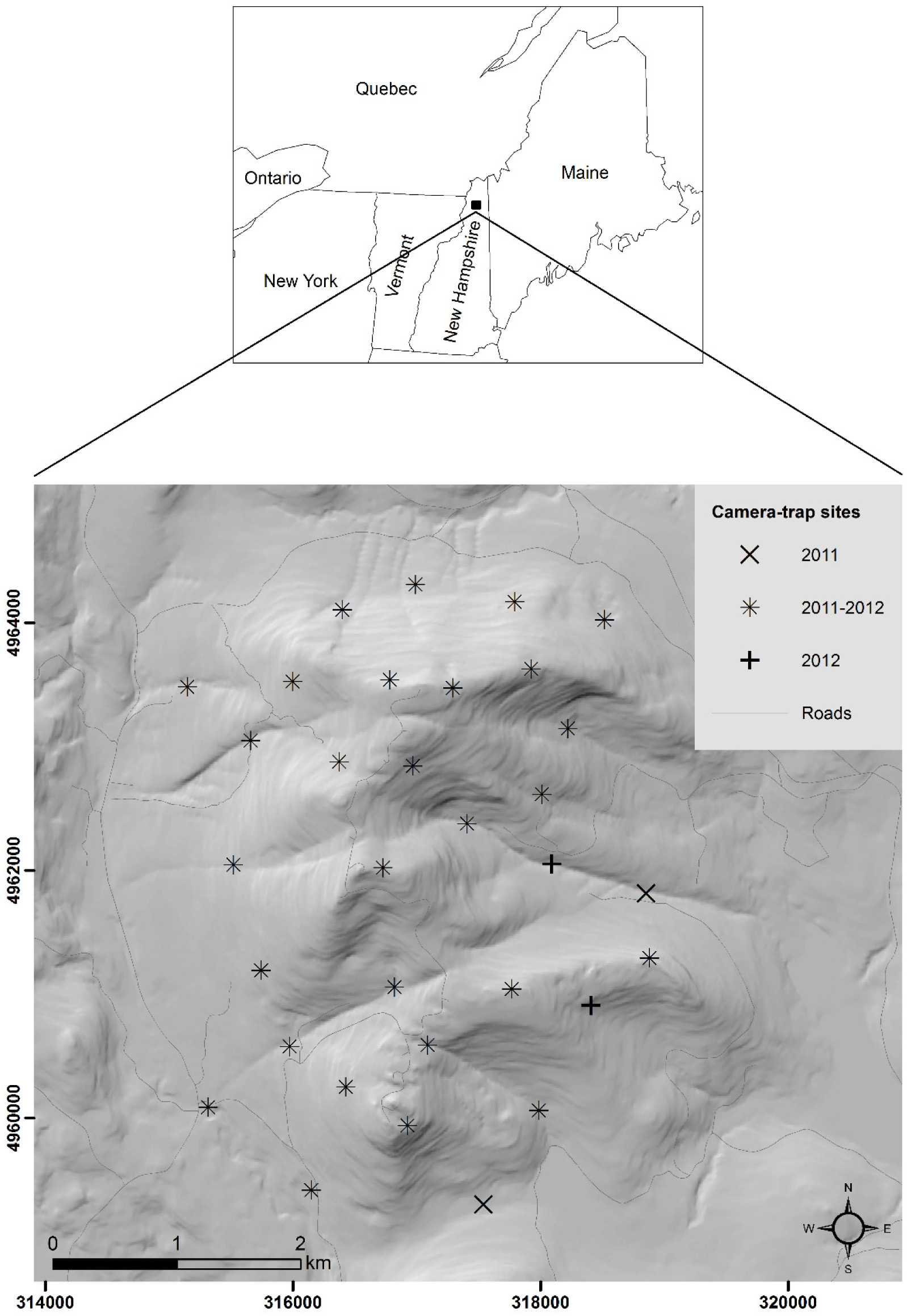
Map of marten study area in northern New Hampshire, USA, reproduced with some modification from Sirén *et al.* (2016a).

### INTEGRATED SCR-RSF MODEL

We estimated multi-scale habitat selection of marten using the spatial encounters of individuals at camera traps and the telemetry locations of collared individuals with the integrated SCR-RSF model (Royle *et al.* 2013). A discrete-space representation of the study area was required to allow spatial environmental covariates to influence the space use of individuals (third-order selection) and the distribution of individuals (second-order selection) on the landscape. The discretization of the landscape accommodates a straightforward formulation of the space usage model that links the camera trap and telemetry data under the integrated SCR-RSF model (Royle et al. 2013). Using the notation of Royle *et al.* (2013), we can define the set of coordinates **x**_1_,…, **x***_nG_* for each pixel *g* on the landscape of *nG* pixels and estimate the likelihood of observing the data for individuals in pixels conditional on **s**, the latent centroids of activity (i.e., home range centers) for individuals, and *z*(**x**), the covariate value(s) for all pixels. Importantly, individual use of a pixel is considered a Poisson random variable with an average rate λ(**x**|**s**) for both data likelihoods. While the total probability of observing an individual in a given pixel differs between the camera trap and telemetry data (as a function of the sampling rate and other features of each device), the mechanisms underlying spatial variation are assumed to be the same, allowing parameters to be shared in the joint likelihood (Royle *et al.* 2013).

Our discrete landscape for marten was defined by a 3.75 km buffer of the trapping array composed of 200 m × 200 m pixels for a total of *nG* = 2,709 non-water pixels. The buffer accommodated individual movement in the sampled population and the pixel resolution was small enough to distinguish differences in space usage within home ranges. We summarized the telemetry data as pixel-specific counts, *m*_*ig*_, for each of *i* = 1,2,…,*N*_tel_ collared individuals at each pixel *g*. Spatial encounters at camera traps, *y*_*ijk*_, were defined for each of *i* = 1,2,…,*n* photo-captured individuals at camera trap *j* on survey *k* as binary variables, condensing any clusters of encounters that occurred in a given day to a single “detection” (Siren et al. 2016a). We extracted the spatial covariate values for trap *j* from the pixel within which the trap was located, such that ***z***_*j*_ ≡ *z*(**x**_*j*_). The spatial covariates we considered for influencing habitat selection included the average distance to mixed-coniferous forest and the vector ruggedness measure (VRM; Sappington, Longshore & Thompson 2007) for each pixel; see Sirén *et al.* (2016b) for more information on the remote sensing data and additional justification for the habitat covariates.

Following Royle *et al.* (2013), we modeled the spatial encounters, *y*_*ijk*_, as Bernoulli random variables such that Pr(*y*_*ijk*_ > 0 | **x***_j_*, **s***_i_*) = *p*_*ijk*_, and under a Gaussian hazard model, *p*_*ijk*_ = 1 – exp(–λ*_ijk_*). In this way, λ*_ijk_* represents the average encounter rate for the trap in pixel **x**_*j*_, assuming the latent “use frequency” (i.e., 3^rd^ order selection) is a Poisson random variable. This use frequency is unobservable for the camera trap data due to difficulty in distinguishing independent encounters within a survey but links directly to the use frequency indicated by the telemetry data (Royle *et al.* 2013). Here, we modeled the encounter rate of marten as a function of several components: 1) availability for an individual, given the location of its latent activity center; 2) resource selection of spatial covariate values for the camera trap; 3) year during which the individual was photo-captured; 4) sex of the individual; and 5) a trap-specific behavioral response. The first two components address the crux of the methodological development presented by Royle *et al.* (2013), while the remaining components are specific to the marten study. Our log-linear model was therefore:

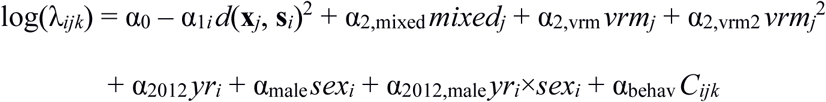

where *d*(**x***_j_*, **s***_i_*) is the Euclidean distance between trap *j* and the latent activity center for individual *i*, and α_1*i*_ = 1/(2σ*_i_*^2^), where σ*_i_* represents the scale parameter of the half-normal distance function. In this way, α_1*i*_*d*(**x***_j_*, **s***_i_*)^2^ quantifies the “availability” of a trap pixel conditional on **s***_i_* for the individual. The spatial covariates were standardized with mean 0 and unit variances, after distance to mixed conifers (*mixed*_*j*_) was square-root transformed and terrain ruggedness (*vrm*_*j*_) was log-transformed. Year and sex were both binary variables indicating whether an individual was encountered in 2012 (*yr*_*i*_ = 1) and its sex was male (*sex*_*i*_ = 1), including a potential interaction. Finally, the behavioral response used *C*_*ijk*_ = 1 for all *k* after the initial encounter of individual *i* at trap *j*, and 0 otherwise. We allowed σ*_i_* to vary according to year and sex:

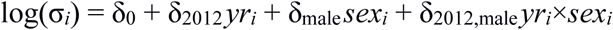

This accounted for potential differences in the scale of movement between sexes and years. We also treated sex as a random variable and estimated φ_male_ = Pr(*sex*_*i*_ = 1) using the model extension described in (Royle *et al.* 2015).

We used the pixel-specific frequencies from the telemetry data to estimate use probabilities under a multinomial distribution such that *m*_*ig*_ ∼ Multinomial(*R*_*i*_, π*ig*), where *R*_*i*_ is the number of telemetry fixes for individual *i* and π*_ig_* is the relative probability of use as defined by

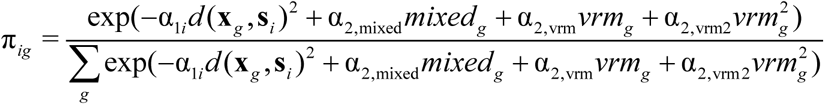

Here, the individual- and pixel-specific usage rate (i.e., the numerator) is formulated with a similar log-linear model as the encounter rate for the spatial encounters at camera traps, with the exception that the rate is not survey specific and is only a function of attributes that vary by location. Another difference is that the usage rate is defined at all pixels (*g*) in the state space, not only the pixels with camera traps. Since the usage rate is a function of availability, the individual rates and relative probabilities are conditional on **s**_*i*_ and variation in the movement scale (σ*_i_*) is defined by the same log-linear model.

We modeled variation in the distribution of activity centers, representing 2^nd^ order resource selection by marten, by specifying an inhomogeneous point process (Borchers & Efford 2008). Here, the expected density in a given pixel *g* was a linear function (on the log scale) of the distance to mixed conifer forest:

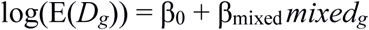

This density model determined the prior probability of an activity center being located in any given pixel according to:

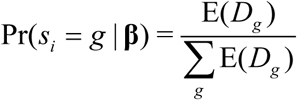

where *s*_*i*_ now represents a pixel ID instead of two-dimensional coordinates. Since the likelihoods of the data for both the spatial encounters and the telemetry fixes are conditional on *s*_*i*_, the marginal likelihood is calculated by integrating over all possible pixel values (i.e., the Poisson-integrated likelihood approach; Borchers & Efford 2008). Note, when considering the photo-captured individuals that were also radio-collared, the conditional likelihoods must be combined before calculating a single marginal likelihood for each individual (Appendix S1). Royle *et al.* (2013) used data from different sets of individuals and assumed the data likelihoods were independent. The consequences of accommodating non-independent data include more precise estimation of activity centers for the captured individuals with collars and reduction of the effective sample size of individuals used to estimate the resource selection functions. Assuming independence between the data when the overlap of individuals is high would result in psuedoreplication problems (e.g., artificially reduced standard errors).

### MODEL COMPARISONS WITH AND WITHOUT TELEMETRY

We compared inferences on multi-scale selection between the integrated SCR-RSF model and a standard SCR model (without telemetry data) using an information theoretic approach followed by parameter estimate comparisons for the relevant coefficients. We used model selection to identify the covariate structures with the best predictive performance for the data at each level of the hierarchical models, starting with movement scale (σ), then encounter rate (λ), and finally density (*D*). A multi-staged approach was used to reduce the total set of candidate models: 1) select among covariates for σ with full covariate structures for λ and *D*; 2) select among covariates for λ using the top covariates for σ and full structure for *D*; and, 3) select among covariates for *D* using the top covariates for σ and λ. Candidate models for the encounter rate always included the behavioral effect (α_behav_) given previous findings (Sirén *et al.* 2016a). This resulted in 5 candidate models for movement scale, 30 candidate models for encounter rate, and 2 candidate models for density. Our main focus was on parameter comparisons for **α_2_** (third-order selection) and **β** (second-order selection), both in terms of best supported model structures and differences in effect size and precision of the estimates, where applicable.

All models were fit using maximum likelihood methods with the oSCR package (Sutherland, Royle & Linden 2016) in R (R Core Team 2016). The oSCR package is an open-source alternative to secr (Efford 2016) for fitting certain spatial capture-recapture models using maximum likelihood. We integrated the R code from Royle *et al.* (2013) into oSCR and included an option for adjusting the marginal likelihood calculations to accommodate non-independent data. The candidate models were ranked using AIC and, to improve clarity for encounter rate comparisons, we removed from our model selection tables those variables with 85% confidence intervals that included zero (Arnold 2010). Note, AIC cannot be directly compared between the standard SCR and integrated SCR-RSF model types given differences in the data likelihoods; we focus on the relative rankings of model structure. We used the top-ranked models to map the predicted encounter probability and expected density across the study area, illustrating variation in resource selection at each scale.

## Results

The winter camera trapping resulted in 13 individuals (6 F; 7 M) captured 121 times in 2011 and 15 individuals (6 F; 9 M) captured 86 times in 2012. Across years, the number of spatial encounters at unique trap pixels per individual ranged 1–10, with 50% having ≤ 2 and 25% having ≥ 5 spatial encounters. The number of radio-collared individuals was 6 (2 F; 4 M) in 2011 and 8 (2 F; 6 M) in 2012, with a total of 147 and 218 telemetry locations, respectively, collected during each winter. Across years, the number of telemetry locations at unique pixels per collared individual ranged 11–27 (median = 17.5). The proportion of encounters at camera traps attributed to radio-collared individuals was 0.73 in 2011 and 0.71 in 2012.

Model selection results indicated that an increased complexity in model structure was supported by the integrated SCR-RSF model with telemetry data compared to the standard SCR model without telemetry data (Tables 1–3). The top model structure for movement scale (σ) included no covariates (i.e., null; AICwt = 0.42) in the absence of telemetry and a sex × year interaction (AICwt = 0.99) with telemetry (Table 1). Sex was also included as a covariate in the top model for encounter rate (λ) without telemetry (Table 2), though model selection uncertainty suggested it was a marginal predictor at best (AICwt = 0.52); none of the spatial covariates were important predictors for encounter rate in the standard SCR model. The integrated SCR-RSF model supported variables corresponding to third-order resource selection (distance to mixed conifer forest and terrain ruggedness) in the top model structures for encounter rate (Table 2). Regardless of telemetry integration, variation in second-order resource selection was supported as distance to mixed conifers was considered an important predictor for density in both the standard SCR and integrated SCR-RSF models (Table 3).

**Table 1.**
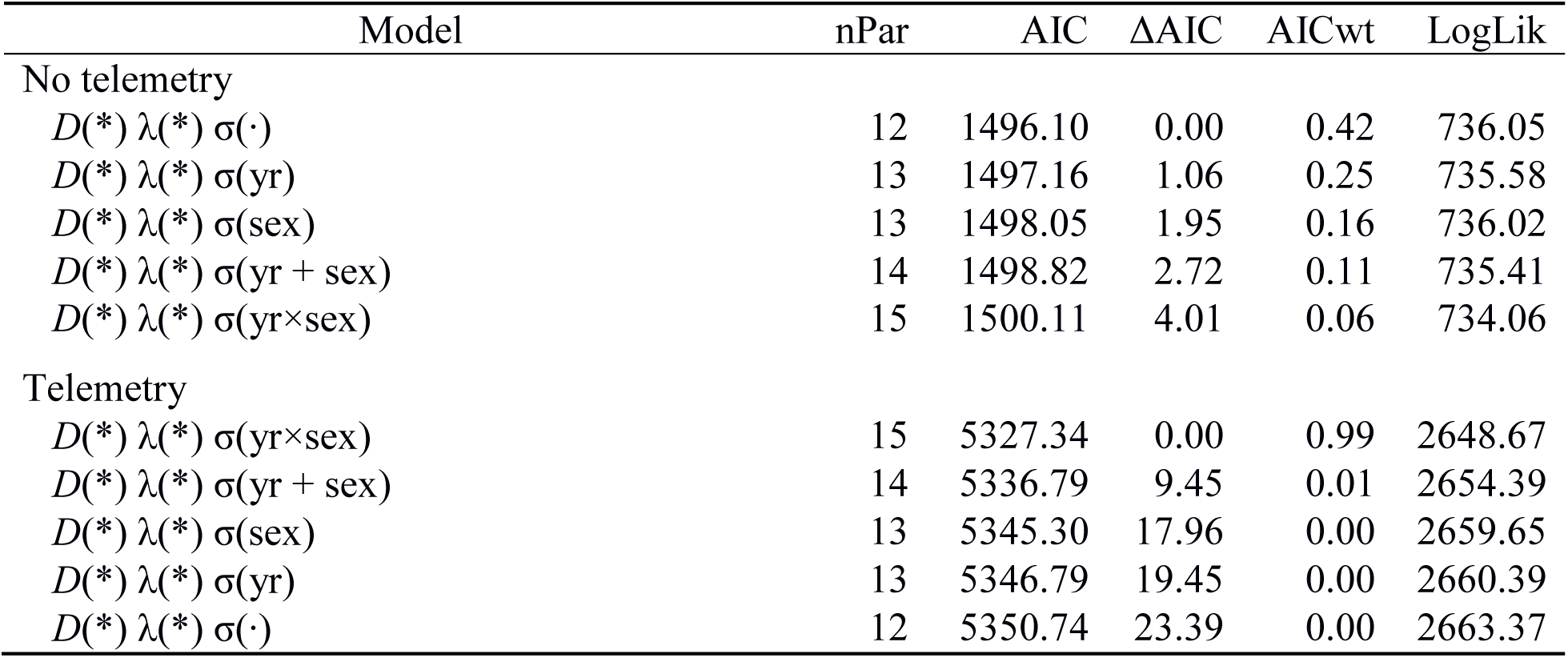
Model selection results for covariates influencing marten movement scale (σ*_i_*) using fully parameterized models for density (D*_g_*) and encounter rate (λ*_ijk_*) both without telemetry and with telemetry integration. Covariates specified here include year (yr) and sex; models with interactions or quadratic terms always included the linear terms.

| Model | nPar | AIC | ΔAIC | AICwt | LogLik |
| --- | --- | --- | --- | --- | --- |
| No telemetry |  |  |  |  |  |
| $D(*) \lambda(*) \sigma(\cdot)$ | 12 | 1496.10 | 0.00 | 0.42 | 736.05 |
| $D(*) \lambda(*) \sigma(\text{yr})$ | 13 | 1497.16 | 1.06 | 0.25 | 735.58 |
| $D(*) \lambda(*) \sigma(\text{sex})$ | 13 | 1498.05 | 1.95 | 0.16 | 736.02 |
| $D(*) \lambda(*) \sigma(\text{yr} + \text{sex})$ | 14 | 1498.82 | 2.72 | 0.11 | 735.41 |
| $D(*) \lambda(*) \sigma(\text{yr} \times \text{sex})$ | 15 | 1500.11 | 4.01 | 0.06 | 734.06 |
| Telemetry |  |  |  |  |  |
| $D(*) \lambda(*) \sigma(\text{yr} \times \text{sex})$ | 15 | 5327.34 | 0.00 | 0.99 | 2648.67 |
| $D(*) \lambda(*) \sigma(\text{yr} + \text{sex})$ | 14 | 5336.79 | 9.45 | 0.01 | 2654.39 |
| $D(*) \lambda(*) \sigma(\text{sex})$ | 13 | 5345.30 | 17.96 | 0.00 | 2659.65 |
| $D(*) \lambda(*) \sigma(\text{yr})$ | 13 | 5346.79 | 19.45 | 0.00 | 2660.39 |
| $D(*) \lambda(*) \sigma(\cdot)$ | 12 | 5350.74 | 23.39 | 0.00 | 2663.37 |

**Table 2.**
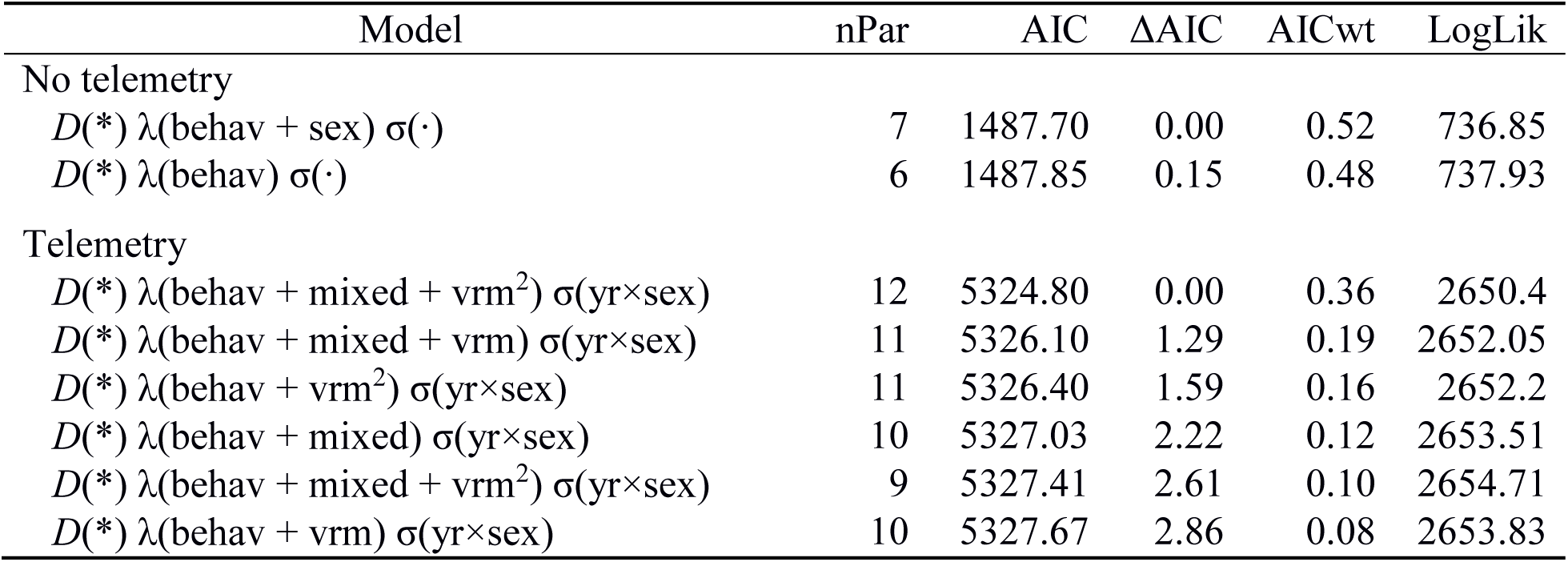
Model selection results for covariates influencing marten encounter rate (λ*_ijk_*) using the top model for movement scale (σ*_i_*) and the fully parameterized model for density (D*_g_*) both without telemetry and with telemetry integration. Covariates specified here include distance to mixed forest (mixed), terrain ruggedness (vrm, vrm^2^), year (yr), sex, and a trap-specific behavioral response (b); models with interactions or quadratic terms always included the linear terms. Models having variables with 85% confidence intervals that included zero are not listed.

**Table 3.**
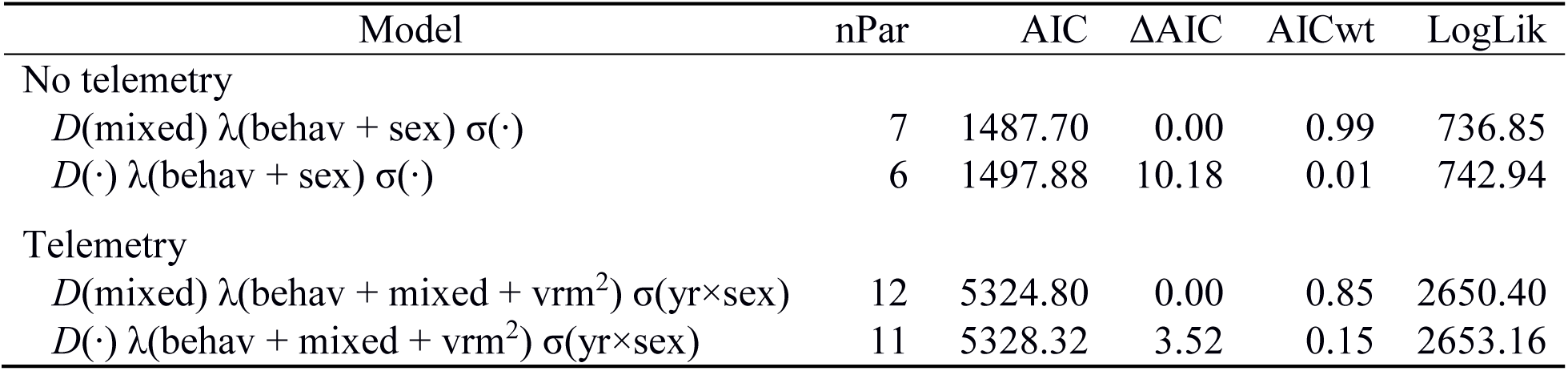
Model selection results for covariates influencing marten density (D*_g_*) using the top models for movement scale (σ*_i_*) and encounter rate (λ*_ijk_*) both without telemetry and with telemetry integration. Covariates specified here include distance to mixed forest (mixed), terrain ruggedness (vrm, vrm^2^), year (yr), sex, and a trap-specific behavioral response (b); models with interactions or quadratic terms always included the linear terms.

Differences in parameter estimates for the top-ranked models further indicated how telemetry integration modified inferences on multiscale resource selection (Table 4). Population density decreased with increasing distance to mixed conifers for both top-ranked models and the effect size was larger without telemetry (β_mixed_ = –2.09 [SE: 0.792]) than with telemetry (β_mixed_ =–0.79 [0.397]). This reduction in effect size for density variation was due to the integrated SCR-RSF model attributing additional variation in the observed encounters to differences in encounter rate, including a decrease with increased distance to mixed conifers (α_2,mixed_ = –0.11 [SE: 0.057]). Encounter rate also appeared to vary by terrain ruggedness, with a positive quadratic relationship suggesting increased space use of terrain that was flat or extremely rugged, and lower use at moderate ruggedness (α2,_vrm_ = –0.11 [SE: 0.055]; α_2,vrm2_ = 0.06 [SE: 0.034]). While maps of expected density illustrated similar spatial patterns of 2^nd^ order resource selection between the top-ranked models (Figure 2a,b), the standard SCR model with no telemetry data exhibited greater variation consistent with the larger estimate for β_mixed_. We also mapped the predicted probability of encounter when *d*(**x***_j_*, **s***_i_*)^2^ = 0 to illustrate the spatial variation in 3^rd^ order selection as indicated by the top-ranked integrated SCR-RSF model (Figure 2c). The overall predicted marten density (#/km^2^) for 2010–2011 was slightly lower for the standard SCR model (0.39/km^2^ [95% CI: 0.29–0.56/ km^2^) than the integrated SCR-RSF model (0.43/km^2^ [95% CI: 0.32–0.61/ km^2^).

**Figure 2.**
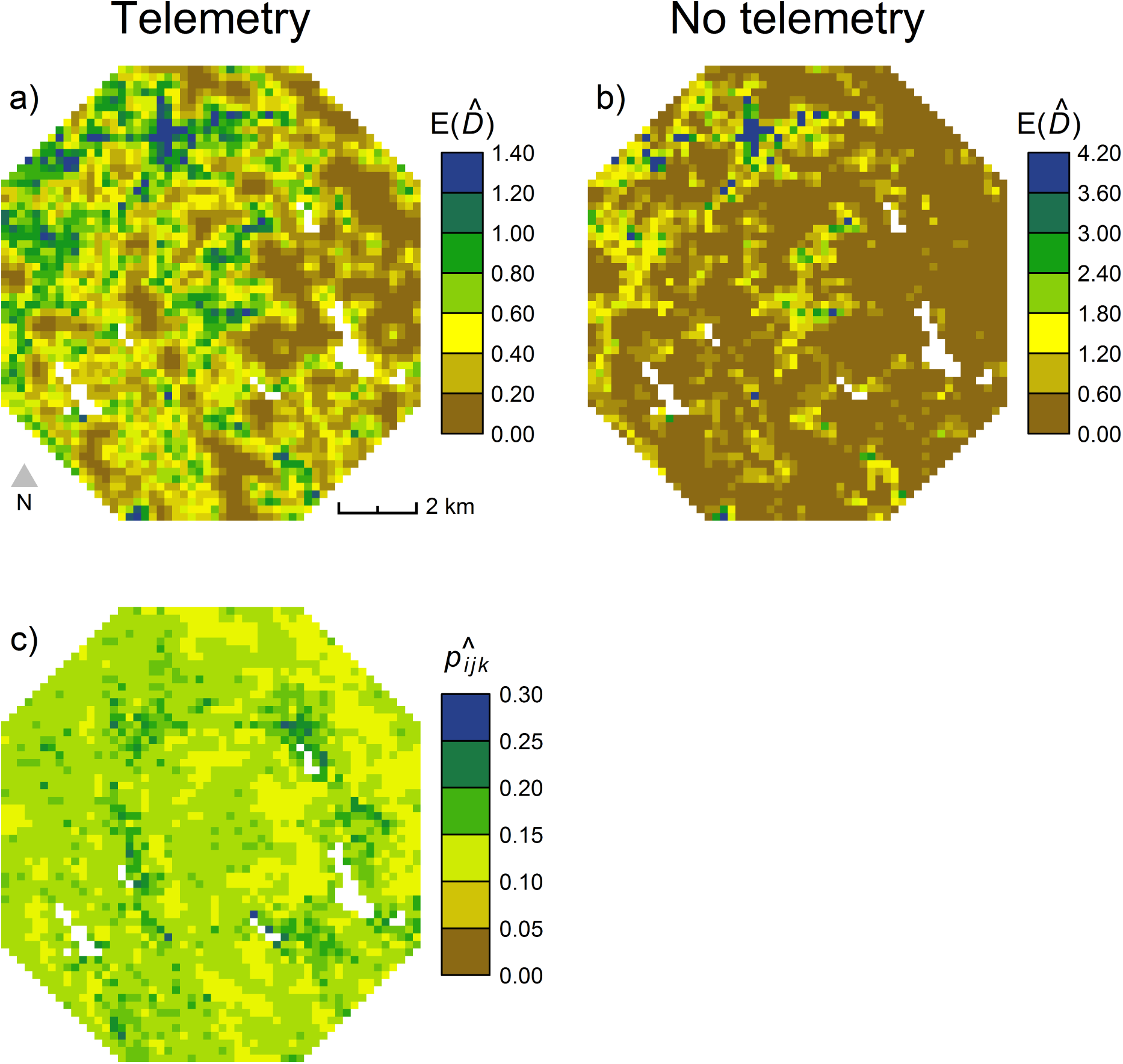
Predicted density of marten (#/km^2^) under the integrated SCR-RSF model with telemetry (a) and a standard SCR model with no telemetry (b), illustrating 2^nd^ order resource selection. In addition, we map the predicted encounter probability for the integrated model (c), illustrating 3^rd^ order resource selection. Blank interior pixels represent water.

**Table 4.** Maximum likelihood estimates (with standard errors) from the top-ranked spatial capture-recapture models.

| Parameter | No telemetry | Telemetry |
| --- | --- | --- |
| Movement scale: $\log(\sigma_i)$ | | |
| $\delta_0$ | -0.23 (0.063) | -0.39 (0.055) |
| $\delta_{2012}$ | — | -0.37 (0.083) |
| $\delta_{\text{male}}$ | — | 0.07 (0.066) |
| $\delta_{2012,\text{male}}$ | — | 0.32 (0.097) |
| Encounter rate: $\log(\lambda_{ijk})$ | | |
| $\alpha_0$ | -2.54 (0.228) | -2.11 (0.141) |
| $\alpha_{2,\text{mixed}}$ | — | -0.11 (0.057) |
| $\alpha_{2,\text{vrm}}$ | — | -0.11 (0.055) |
| $\alpha_{2,\text{vrm}2}$ | — | 0.06 (0.034) |
| $\alpha_{2012}$ | — | — |
| $\alpha_{\text{male}}$ | 0.29 (0.201) | — |
| $\alpha_{2012,\text{male}}$ | — | — |
| $\alpha_{\text{behav}}$ | 1.33 (0.174) | 1.19 (0.151) |
| Density: $\log(E(D_g))$ | | |
| $\beta_0$ | -5.58 (0.869) | -4.32 (0.311) |
| $\beta_{\text{mixed}}$ | -2.09 (0.792) | -0.79 (0.397) |

The top-ranked models also differed with regards to movement scale (σ) variation and the predicted probabilities of activity center locations for collared individuals. The standard SCR model did not support any variation in movement scale, with an average σ = exp(δ_0_) = 0.79 km (95% CI: 0.70–0.90 km), while the integrated SCR-RSF model suggested an interaction between sex and year such that female marten in 2012 had a lower movement scale than all other individuals (Table 4). The precision of the activity center predictions was significantly increased for collared individuals when telemetry data were integrated (Figure 3), underscoring the value of using a non-independent likelihood.

**Figure 3.**
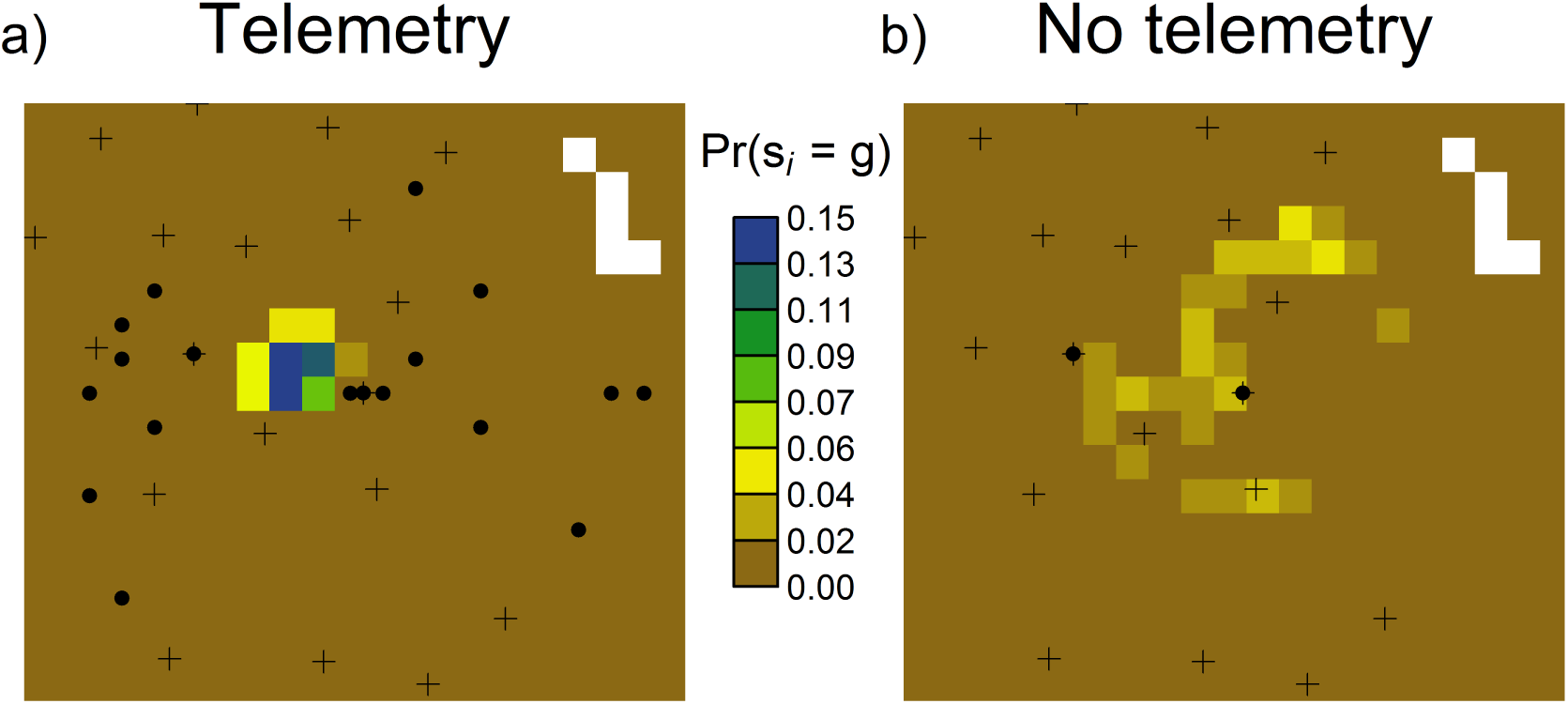
Probability densities of the activity center *s*_*i*_ in pixel *g* for one radio-collared male marten in 2012 from the integrated SCR-RSF model with telemetry (a) and a standard SCR model with no telemetry (b). Crosses represent camera traps and black dots represent spatial encounters within a 200 m pixel recorded by cameras and/or telemetry fixes.

## Discussion

Integrated modeling methods allow ecologists to bring together multiple data sources to expand the scope of potential inferences for and enhance understanding of complex ecological systems. The recent growth of these techniques coincides with the development of hierarchical models (Gimenez *et al.* 2014), which posit data generation as a combination of observation processes and latent ecological processes (Royle & Dorazio 2008). When multiple sources of data provide information on the same ecological processes of interest, joint data likelihoods can be defined to explicitly link diverse observations and improve estimation of model parameters describing relevant phenomena. The result is an increased ability for ecologists to join empirical data to ecological theory using robust quantitative approaches.

The integrated SCR-RSF model developed by Royle *et al.* (2013) connects observed spatial encounters of animals to established theory on multi-scale resource selection (Johnson 1980). By combining data from devices that differ in their ability to target variation at specific scales (i.e., individual vs. population level), the model is better able to simultaneously describe both scales. Our application of the integrated SCR-RSF model allowed for more complex covariate relationships with parameters related to individual movement and multi-scale resource selection of marten than possible under standard SCR model fitting. In the absence of telemetry data, spatial variation in third-order selection (i.e., through encounter rate) could not be detected with the camera trap observations and instead, the standard SCR model suggested more variation in second-order selection. Both model types indicated that expected marten density decreased with increasing distance to mixed conifers. So while the camera trap data alone were sufficient for identifying an existing density relationship, the slope of this relationship was potentially biased due to unmodeled heterogeneity in encounter rate. The integrated SCR-RSF model indicated that areas of the landscape with relatively short distances to mixed conifer forest had more marten, and those marten spent more time in these locations. Additional model complexity supported by our telemetry integration included an interaction between sex and year on movement scale (σ), with female marten in 2012 being encountered at shorter distances from their activity centers than individuals of either sex in other years. The standard SCR model had little power to detect variation in σ according to locations of spatial encounters at camera traps alone, likely due to small sample sizes. Similarly, the small number and strategic placement (i.e., to increase encounters) of camera traps made it difficult to detect variation in encounter rate according to spatial environmental variables without the additional data provided by telemetry.

Telemetry has been used in multiple ways as an auxiliary data source to improve inferences from population size estimation, including both ad-hoc adjustments (Soisalo & Cavalcanti 2006) and more recent joint modeling techniques (Ivan, White & Shenk 2013; Bird *et* al. 2014). Spatially-explicit applications have generally focused on the contribution of telemetry data to informing the scale of individual movement in SCR models for the purposes of improving the precision and/or identifiability of parameter estimates (Sollmann *et al.* 2013a; Sollmann *et al.* 2013b). Even as few as *N*_tel_ = 3 collared individuals with an adequate collection of telemetry locations can greatly increase the precision of the σ estimate (Royle *et al.* 2013), though such small sample sizes will limit the exploration of potentially important relationships with individual attributes (e.g., sex; Sollmann *et al.* 2013a). Here we had some flexibility with *N*_tel_ = 14 collared marten and were able to model more variation in σ, though at the expense of a significant improvement in precision (Table 4). Most importantly, we were better able to model variation in encounter rate as it related to individual space use due to the relatively large collection of telemetry locations.

Proffitt *et al.* (2015) is the only other application we could find where telemetry-informed resource selection was integrated into an SCR framework in the vein of Royle *et al.* (2013). Their analysis involved a two-stage approach where 30+ years of radiotelemetry data on mountain lions (*Puma concolor*) were used to estimate and validate an RSF that corresponded to second-order selection (Robinson *et al.* 2015); spatial predictions from the RSF were then specified as a density covariate in an SCR model fitted to search encounter data collected during a 4-month sampling period. The RSF covariate appeared to improve the accuracy and precision of density estimates (Proffitt *et al.* 2015), yet, without integrating the telemetry data into the likelihood directly, their analysis could not incorporate information on third-order selection or benefit from increased precision in σ estimates. The latter may have provided a significant improvement due to the small number of spatial encounters and could have alternatively been achieved in their Bayesian modeling approach by using an informed prior for σ calculated from the 18,000+ telemetry locations on 85 individuals (Proffitt *et al.* 2015). The individuals in the telemetry data need to be representative of the focal population being exposed to capture and the degree to which this is true will dictate how many model parameter estimates can reasonably be shared between the data.

The marten study consisted of individuals that overlapped entirely between the telemetry and camera-trap data, as all radio-collared individuals were also photo captured. While this obviously addressed concerns about collared individuals being representative of the focal population targeted by camera traps, it required a modification to the joint likelihood (Appendix S1) originally constructed under an assumption of data independence (Royle *et al.* 2013). The primary benefit to our joint likelihood came from the increased precision of predicted activity centers for collared individuals (Figure 3), which theoretically should have improved estimation of the inhomogeneous point process model for density and the conditional probabilities of encounter. The resulting inferences were mostly similar to the separate analyses previously described by Sirén *et al.* (2016a) and Sirén *et al.* (2016b) regarding average density and multi-scale resource selection, respectively, of American marten in alpine forests of New England. Sirén *et al.* (2016b) used a “design III” approach to fitting an RSF (Manly *et al.* 2002) and found that regenerating forest was the most important factor influencing second-order resource selection by marten, as individuals tended to avoid it on the landscape. Here, our inhomogeneous point process was akin to a “design II” approach (Manly *et al.* 2002) and we used proximity metrics (e.g., distance to feature) instead of compositional metrics for the spatial habitat covariates; therefore, some differences were to be expected. In other landscapes of eastern North America, marten have similarly exhibited second-and third-order selection for older-aged mixedwood forest (Potvin, Bélanger & Lowell 2000; Cheveau *et al.* 2013). In the present analysis, density estimation was simultaneously modulated by both second-order and third-order resource selection, which were found to be important predictors of variation in the observed spatial encounters for both camera traps and telemetry locations.

In summary, the integrated SCR-RSF model addresses concerns regarding heterogeneity in capture due to individual space use which can otherwise generate bias in the estimation of density using spatial capture-recapture models (Royle *et al.* 2013). The increased popularity in using SCR to estimate density of rare, wide-ranging species (e.g., carnivores) will result in many sparse datasets that are unlikely to support complex encounter models (Sollmann *et al.* 2013a). Adding several individuals with VHF or GPS collars to provide an auxiliary source of movement information can increase the accuracy and precision of inferences from spatial encounter designs, particularly when species are selecting resources at multiple scales. Using the modified likelihood as made available in oSCR (Sutherland, Royle & Linden 2016) will allow researchers with data sets containing heavy overlap of individuals to fit integrated SCR-RSF models that can accommodate the lack of independence and improve parameter estimation. Integrated modeling approaches allow ecologists to make inferences with the best available information and improve our understanding of ecological systems and our ability to develop effective conservation and management strategies.

## Acknowledgments

Funding for this project was provided through New Hampshire State Wildlife Grant T2-1-R in cooperation with the United States Fish and Wildlife Service. Additional support was provided by the Department of Interior, Northeast Climate Science Center. We would like to thank Wagner Forest Management, American Forest Management, and lessee Brookfield Renewable Power for providing access to the study area. The New Hampshire Fish and Game Department, United States Fish and Wildlife Service, and Stantec Environmental Consultants provided invaluable assistance, equipment, and lodging. W. Staats, J. Kilborn, A. Timmins, H. Jones, and Z. Smith provided invaluable field assistance.

## Data accessibility

Data will be archived with Dryad Digital Repository.

## Appendix S1. Likelihood for integrated SCR-RSF model with non-independence

Royle et al. (2013; Supplement 1) described the full likelihood for the integrated SCR-RSF model assuming that the capture data and telemetry data were independent. To accommodate the marten study where all collared individuals were also photo-captured, the conditional likelihoods need to be combined before calculating the marginal likelihood for each individual. We do not describe all components of the likelihoods here as they are fully described in Royle et al. (2013; Supplement 1); we instead emphasize the main differences between the independent and non-independent formulations. In addition, our model description here is simplified to match that of Royle et al. (2013), removing some details specific to the marten study.

When the datasets are independent, the total likelihood for the integrated SCR-RSF model is the product of the likelihoods for the capture data (SCR) and the telemetry data (RSF):

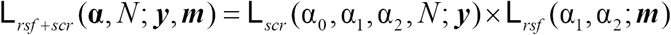

Here the **α** parameters correspond to variation in the encounter rate (for spatial encounters, ***y***) and usage rate (for telemetry locations, ***m***), while population size (*N*) is only estimated from the SCR model. Note that α_0_ only appears in the SCR likelihood as it corresponds to the baseline encounter rate and does not involve spatial variation. The other parameters represent availability (α_1_), conditional on the latent activity centers (**s**), and resource selection (α_2_).

The conditional-on-**s** likelihoods, here represented as *f*(data | **s**, parameters), differ according to the observation models for the capture data and the telemetry data. The marginal distributions, here represented as *f*(data | parameters), for each individual *i* are calculated by integrating the conditional-on-**s** likelihoods over the possible locations for the individual activity centers, **s***_i_*:

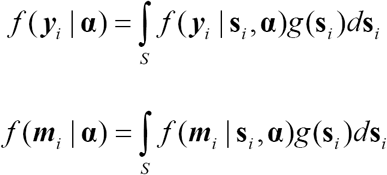

Here, *S* represents the continuous state space over which the integral is calculated and g(**s***_i_*) = 1/||*S*|| when density does not vary, indicating a homogeneous point process for activity centers (note: we used a discrete state space and an inhomogeneous point process (Borchers & Efford 2008) for the marten study). The likelihoods for all observed individuals in the data (*n* for captures, *N*_tel_ for telemetry) are then the products of the individual likelihoods:

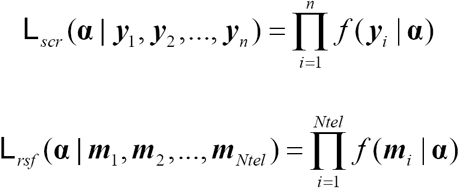

As described earlier, the product of these two data likelihoods then provides the total likelihood. Royle et al. (2013; Supplement) provide a more complete description of the likelihoods, including the important contribution of unobserved individuals for estimating *N* from the capture data. We do not highlight the components of the likelihood that remain the same regardless of independence between datasets.

In applications where the data sources are not independent (including the marten study), the individual conditional-on-**s** likelihoods for each data source must be combined before computing the marginal likelihood for each individual. When all collared individuals are also captured, the marginal likelihood for collared individuals is as follows:

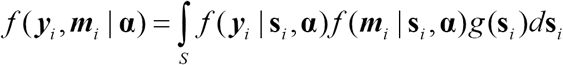

The marginal likelihood for the *n* – *N*_tel_ individuals that were captured but not collared does not change. The joint likelihood for the observations, assuming collared individuals are sorted first, is then:

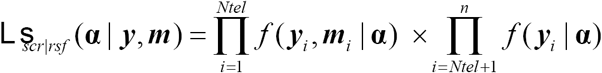

In this way, the likelihood for the telemetry data becomes embedded in the likelihood for the capture data for those collared individuals that were also captured. If there were also some collared individuals that were never captured, the joint likelihood would involve the product of the two components above and a third representing the marginal distribution of the telemetry data. Thus, depending on how individuals overlap between data sets, there are three potential marginal distributions to calculate for the observed individuals: 1) captures only; 2) collars only; and 3) captures and collars. Assuming independence, only #1 and #2 are used in the joint likelihood. The marten study here consisted of #1 and #3. The integrated likelihood in oSCR (Sutherland, Royle & Linden 2016) currently supports model fitting to data consisting of #1 and combinations involving #1 and #2, and #1 and #3.

